# A surprising link between cognitive maps, successor-relation based reinforcement learning, and BTSP

**DOI:** 10.1101/2025.04.22.650046

**Authors:** Yukun Yang, Christoph Stöckl, Wolfgang Maass

## Abstract

Recent recordings from the hippocampus of the human brain suggest that after a few presentations of sequences of unrelated natural images, correlations between emergent neural codes encode the sequential structure of these sequences. We show that this learning process, which is consistent with experimental data on BTSP (Behavioral Time Scale Synaptic Plasticity), creates a cognitive map that enables online-generation of plans for moving to any given goal, both in spatial environments and in abstract graphs. Furthermore, the resulting neural circuits and plasticity rules provide a biologically plausible implementation of Successor Relation based Reinforcement Learning. In addition, this brain-derived approach for learning cognitive maps provides a blueprint for implementing autonomous learning through on-chip plasticity in energy-efficient neuromorphic hardware.

## 1 Introduction

Planning and problem solving are fundamental higher cognitive functions of the brain (Summerfield, 2022). But according to the review (Mattar and Lengyel, 2022) it remains largely unknown how the brain achieves that. Planning is defined in this review as *the process of selecting an action or sequence of actions in terms of the desirability of their outcomes*. Importantly, this desirability may depend on distant outcomes in the future, hence some form of look-ahead is required for efficient planning. Many planning tasks and problem solving tasks can be formulated on an abstract level as tasks to find a shortest path from a given start to a given goal node in some graph, see the first chapters on planning and problem solving in the standard AI textbook (Russell and Norvig, 2020). This graph is in general not planar, and its nodes may not necessarily represent locations in some physical environment, but rather encode combinations of external and/or internal states in a planning environment.

Numerous experimental data from the rodent and human brain have shown that cognitive maps are likely to support navigation in spatial environments and also in 2-dimensional concept spaces (Behrens et al., 2018; Bottini and Doeller, 2020). A first model for learning cognitive maps that enable efficient online planning has been proposed in (Stöckl et al., 2024). This approach is based on learning embeddings of observations and actions into a common high-D (high dimensional) space. Learning of these embeddings is driven by the goal to predict the next observation based on the current observation and the current action. We show here that functionally useful cognitive maps also arise in the context of quite different learning processes: The formation of neural codes for sequences of observations, i.e., of memory traces for episodic memories. Their emergence after exposure to just a few repetitions of sequences of natural images was recently shown through electrode recordings from the human brain (John et al., 2025; Tacikowski et al., 2024).

It currently remains unknown which synaptic plasticity mechanism creates memory traces for episodic memories in the human hippocampus. But substantial experimental evidence shows that episodic memories, or more generally conjunctive codes for diverse items, are formed in area CA1 of the rodent by BTSP (Behavioral Time Scale Synaptic Plasticity) (Bittner et al., 2017). Importantly, BTSP was shown to open the gate for synaptic plasticity for several seconds, and therefore acts on the time scale of image sequences of the experiments of (John et al., 2025; Tacikowski et al., 2024). Furthermore, it was shown that BTSP is already induced by a single trial, and instantaneously expressed. A simple model for BTSP that can reproduce many of the underlying experimental data was recently proposed in (Wu and Maass, 2025). We show here that this model for synaptic plasticity can also reproduce the experimental data on fast learning of sequences from (John et al., 2025; Tacikowski et al., 2024). In particular, it produces correlated neural codes for observations that co-occur in some sequences (episodes).

In order to plan one also needs to link observations with actions. In fact, this interlinking has been proposed to be fundamental for the emergence of brain function (Buzsáki, 2019). We therefore interlink in the proposed model for planning the previously sketched learning process, whereby observations become represented by neural codes for sequences in which they occur, with another simple local rule for synaptic plasticity that relates these neural codes for observations with neurons that initiate actions that could be causal for inducing these observations. The resulting activation of action neurons through observations of their outcome is actually closely related to experimental data on mirror neurons (Bonini et al., 2022).

We demonstrate that the resulting simple learning model, to which we refer as correlation based Cognitive Map Learner (CCML), provides online planning capability for solving complex tasks, such as navigation to arbitrary goals in a given graph. Importantly, the powerful functional capabilities of the CCML can be explained through a rigorous theory. In fact, the CCML can be seen as neural implementation for the arguably most powerful algorithmic model for learning flexible goal-directed behavior: Reinforcement Learning (RL) based on the Successor Relation (SR), originally proposed by (Dayan, 1993). SR-based RL has since then be argued to provide from the theoretical perspective the most plausible model for human RL (Momennejad et al., 2017). But a biologically plausible implementation of SR-RL in neural circuits of the brain has been missing. We show here that planning by the CCML provides a neural implementation model for SR-based RL.

CCMLs share with Transformers (Vaswani et al., 2017) that they do not require a teacher for learning, and that the outcome of learning is an embedding of external tokens into a high-dimensional internal model. But in contrast to Transformers, the CCML neither requires deep learning nor large amounts of data: Its learning can be implemented through local synaptic plasticity, and it only needs a moderate amount of exploration. In fact, an implementation of a CCML in energy efficient neuromorphic hardware is even simpler than an implementation of the prediction-learning based CML of (Stöckl et al., 2024). In particular, the CCML appears to be well-suited for enabling autonomous learning of goal-directed behavior through on-chip synaptic plasticity of neuromorphic hardware.

## 2 Results

Learning of a cognitive map in the CCML (correlation-based cognitive map learner) is based, like for the CML of (Stöckl et al., 2024), on learning an embedding **Q** of observations **o** into some high-D space. We often refer to resulting high-D vectors **Q**(**o**) as states. Observations can consist of sensory inputs and/or internal state representations. The goal of this embedding is that structural relations between the embedded points **Q**(**o**) encode information about how the corresponding observations are related to each other by actions, or chains of actions. The CCML uses correlations between these state representations to encode how easy or difficult it is to reach from one to the other through a sequence of actions. The idea for the definition of **Q** in the CCML is provided by the experimental data of (John et al., 2025; Tacikowski et al., 2024). The correlation between internal representations of two different images is according to their data closely related to the number of sequences in which both of them occur.

We consider the simplest possible model for these experimental data: Neural codes for embedded observations (images) consist of sparse binary vectors, where a “1” denotes firing of a corresponding neuron for the image. Thus each neuron in the population corresponds to one dimension in the high-D coding space, and is dedicated to one of the experienced sequences (episodes): It fires for all observations that occur in this sequence and for no other observations, see Fig. 1A, B. Then the dot-product (**Q**(**o**) · **Q**(**o**′)) between the embeddings of two observations **o** and **o**′ is equal to the number of sequences in which both observations occur. The dimension *N*_*s*_ of the high-D coding space is equal to the number of sequences that were presented.

**Fig. 1:**
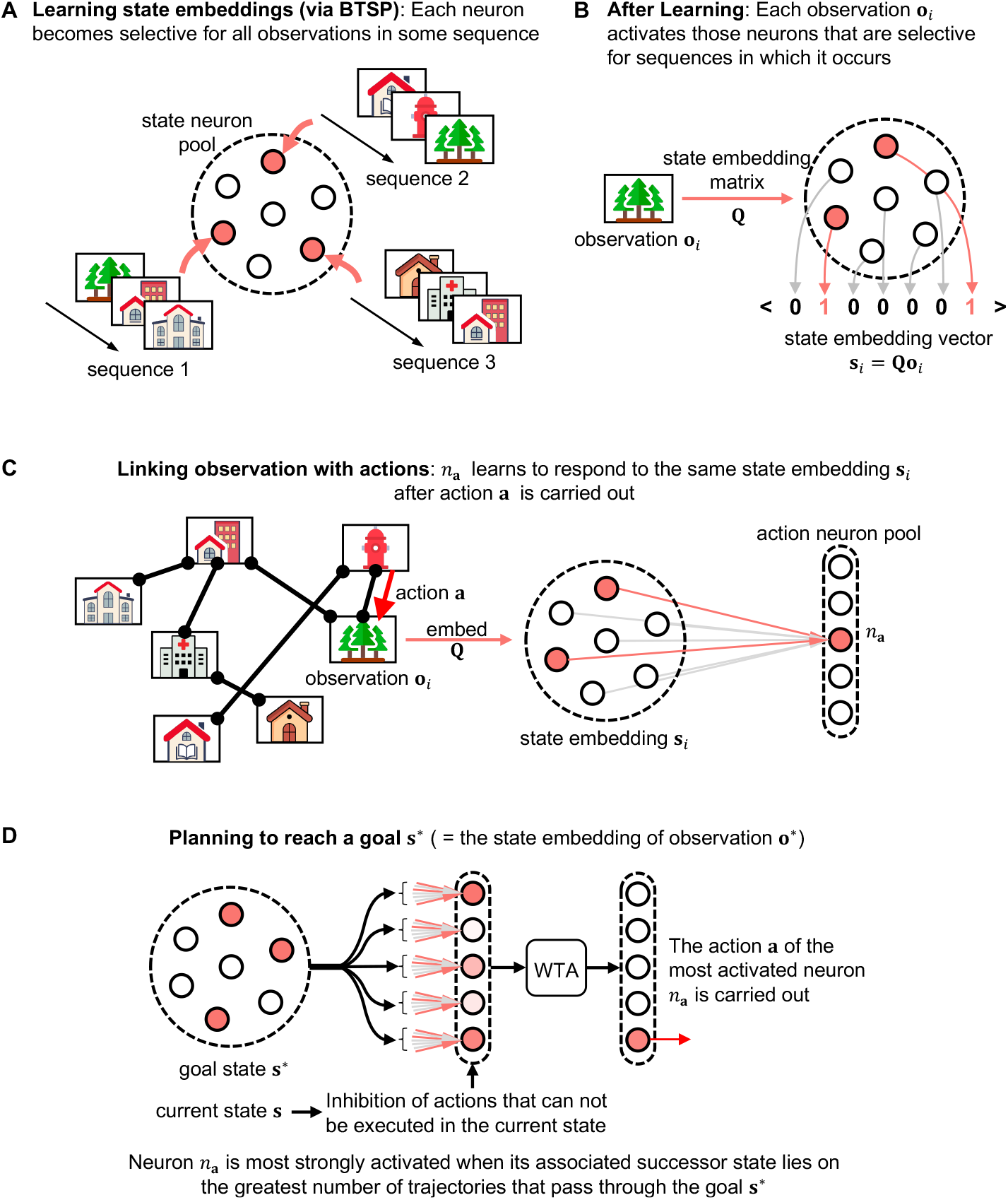
Architecture of the CCML. **(A)**. Illustration of the embedding **Q** of observations **o** into a high-D space, where one neuron is dedicated to each sequence of observations that is presented, and fires for all observations **o** in this sequence. **(B)**. Illustration of the resulting binary code for embedded experiences **Q**(**o**). **(C)**. Architecture for learning to associate the next observation to the action **a** that caused it. 1-shot Hebbian learning with binary synaptic weights suffices. **(D)**. Goal-directed action selection in the resulting cognitive map can be implemented through a WTA (Winner-Take-All) competition, applied to all actions **a** that can be executed in the current state.

In the behavioral experiments of (John et al., 2025; Tacikowski et al., 2024) this ensemble of sequences was the set of sequences that were repeatedly presented to the human subjects. Our model is also compatible with an online scenario where the number *N*_*s*_ of these sequences grows over time, yielding neural coding vectors that grow in length when additional neurons are recruited for representing new sequences. Links between our model for learning the embedding **Q** and experimental data on emergent sequence representations in the human hippocampus are exhibited in Fig. 2.

**Fig. 2:**
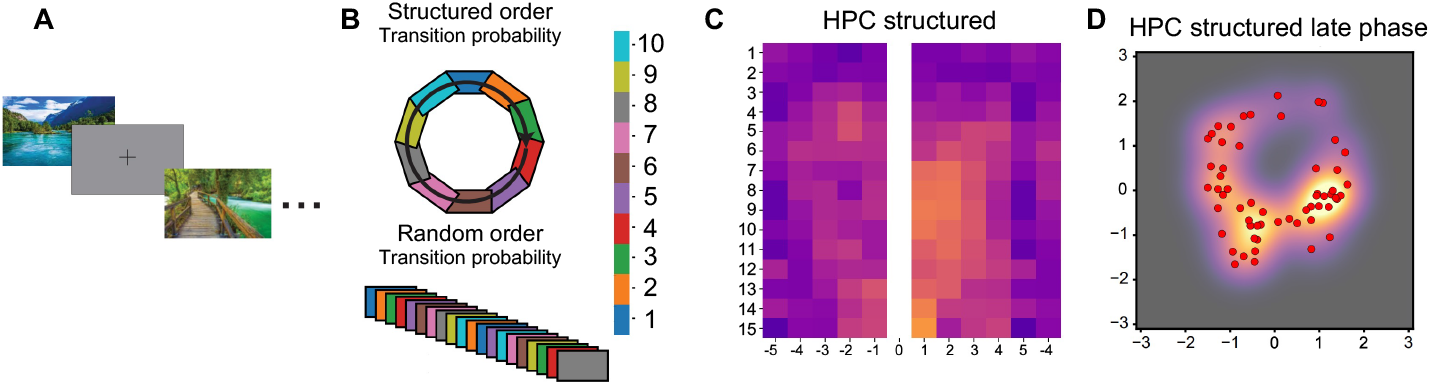
Links between the embedding Q of the CCML and experimental data on emergent representations of sequences in the human hippocampus. (All panels are copied from (John et al., 2025).) **(A)** Setup of their experiment: 10 natural images were repeatedly presented to human subjects, each for 1s with a blank screen for 1s in between. **(B)** In the “structured” version, images were presented in 15 repeated cycles of a fixed order, leading to the results shown in **C** and **D**. By contrast, presenting the images in random order did not produce any patterned results. **(C)** Temporal evolution of the firing response of a neuron (indicated through heat map, lighter color indicates more firing) to images that were presented before and after a specific image (at position 0 in this plot). One sees that already after 3 presentations of the sequences the neuron starts to respond to adjacent images in this sequence. **(D)** A ring-shaped “cognitive map” emerged during later rounds of the cyclic image presentations in a 2D projection of recorded LFP signals from the human HPC (each natural image presentation during the last rounds gives rise to one data point).

In order to enable planning for reaching arbitrary goals, that take here the form of target observations **o**^∗^, one need to learn besides the embedding **Q** also a linear map **W** that maps embeddings **Q**(**o**) of observations **o** onto the actions **a** that preceded it in a sequence. More precisely, we consider a population of linear neurons *n*_*a*_ that vote for different actions **a** and **W** consists of the synaptic weights of these neurons action selection neurons. This map **W** corresponds to the inverse of the action embedding **V** that was learnt in the model of (Stöckl et al., 2024). **W** can be learnt in the case of the CCML by a simple 1-shot Hebbian learning rule: Set the weights of the neuron equal to the embedding **Q**(**o**′) of the next observation **o**′ that results from carrying out action **a**. Note that we assume that each action can only be carried out in a specific state, and produces deterministically a specific observation **o**′. This ensures that the weights of neuron *n*_*a*_ are subject to a single application of the Hebbian 1-shot learning rule, and have therefore the values given by **Q**(**o**′). The CML from (Stöckl et al., 2024) appears to be better suited to navigation in spatial environments where the same action can be applied in many different states (= places).

Surprisingly, for planning a sequence of actions that lead to a target observation **o**^∗^ it suffices for the CCML to present **Q**(**o**^∗^) as synaptic inputs to all neurons *n*_*a*_, and to choose among all actions **a** that can be carried out in the current state that one for which the linear neuron *n*_*a*_ is most strongly activated, i.e., has the highest utility in the terminology of (Stöckl et al., 2024). We show in Fig. 3 that this simple online-planning method is competitive with the optimal offline planning method, the Dijkstra algorithm, with regard to lengths of resulting paths to the goal.

**Fig. 3:**
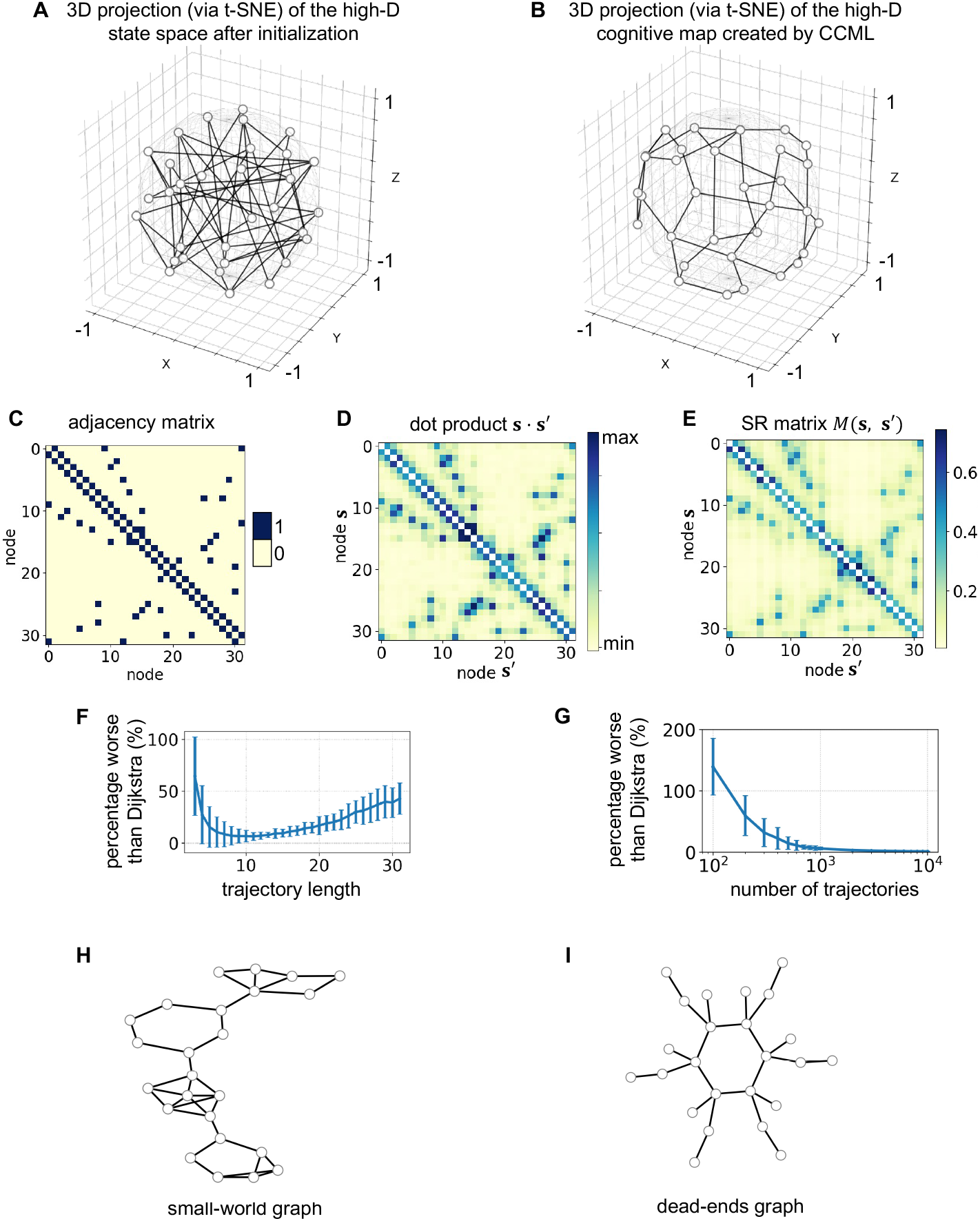
Performance of the CCML for planning in an abstract graphs. **(A)** Initial state embeddings for a random graph. **(B)** Embeddings after CCML learning. **(C)** Graph adjacency matrix. **(D)** Pairwise dot-product heat map of embeddings. **(E)** Successor-representation matrix under a random policy. **(F)** Planning performance (vs. Dijkstra) versus trajectory length for 1000 random walks. Agent takes random walk in the environment, at each step, it truncate a fixed length time window back to its past that generates one trajectory. **(G)** Planning performance versus number of trajectories (length = 10); both **F–G** show mean ± standard deviation over 100 randomly generated graphs. One sees that the model already has near-optimal performance when the agent moves on the graph for 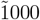 steps. **(H)** Small-world graph: near optimal performance (4.48% worse than Dijkstra averaged over 10 randomly generated small world graphs) is reached with 10000 walks and cutted into trajectories with window length 20; The number of trajectories may be reduced dramatically if the agent switches to goal-directed trajectories after some initial random-walks. **(I)** Dead-end graph: optimal performance is reached with 1000 walks of length 10.

Fig. 3A-B shows the cognitive map before and after learning in 3D (projected from high-D back to 3D via t-SNE, then normalized to unit-length vectors). The CCML agent is trained on 1000 sequences with each containing 5 nodes (The agent moves a very long trajectory length 1004 which we divided into 1000 overlapping trajectories with each length 5). The resulting geometry enables greedy moves toward nodes whose embeddings best correlates with the goal. Detailed performance of CCML under different settings of trajectory length and number of trajectories can be found in 3F-G.

The good performance of this planning method can be rigorously explained by applying the theory of SR-based RL. The SR is a relation *M* (**s, s**′) between states **s** and **s**′ that records the expected number of future discounted visits to state **s**′ during some exploration policy, such as random walks from state **s** (Momennejad et al., 2017). It can be learnt through temporal difference (TD) learning. The dot-product between embeddings **Q**(**o**), **Q**(**o**′) of observations **o** and **o**′ in our model is a closely related quantity: it counts in how many exploration sequences both the observation **o** and the observation **o**′ were encountered. If these exploration sequences are segments of a random walk, then this dot product can be seen as empirical estimate of a SR (See 3C-E. 3D uses the same settings in generating 3B, and 3E uses the discount factor *γ* = 0.8). In SR-based RL one uses then for reaching some arbitrary goal **g** the SR *M* (**s, g**) as value function *V*_**g**_(**s**) for any state **s**. A greedy policy for reaching a goal **g** moves then from any current state **s** to that adjacent state **s**′ for which *M* (**s**′, **g**) is maximal. We refer to the Methods section for details.

In the previously described planning method of the CCML one chooses among currently affordable actions **a** that action for which the linear neuron *n*_*a*_ with synaptic input **Q**(**o**^∗^) produces the largest output. Remember that the synaptic weights of this linear neuron were defined during learning as **Q**(**o**′) for the next observation **o**′ that results from carrying out this action **a**. Hence this action selection of the CCML amounts to choosing that action **a** for which *M* (**s**′, **g**) = (**Q**(**o**′), **Q**(**o**^∗^)) is maximal, where **s**′ = **Q**(**o**′) is the state that results from action **a** and **g** = **Q**(**o**^∗^) is the goal. This proves that the CCML chooses in each state the same action that a SR-based RL agent would choose. We show in Fig. 3 that this simple online planning method performs very well for navigation to arbitrary given goals in undirected abstract graphs.

A closer look shows that one structural difference remains between CCMLs and SR-based RL: If the direction in which sequences of observations are encountered during exploration is not taken into account, as we do in our previously described version of the CCML model, then the dot product (**Q**(**o**) · **Q**(**o**′)) is a metric, in particular it is symmetric. In contrast, the SR is not automatically symmetric. We will show in Fig. 4 that a refined version of the CCML where the direction of exploration trajectories is taken into account is also able to carry out very efficient planning to arbitrary goals in directed graphs. The only difference is that for reaching a goal **g** = **Q**(**o**^∗^) one gives as input to the neurons *n*_*a*_ instead of the binary vector **Q**(**o**^∗^) a reduced binary vector where 1’s remain only at those positions that correspond to trajectories where the next state **s**′ = **Q**(**o**′) after carrying out action **a** comes prior to **Q**(**o**^∗^) or is equal to **Q**(**o**^∗^). Note that experimental data suggest that brains to record in which direction a sequence of observations were encountered. According to (Sugar and Moser, 2019) an episodic memory encodes for each of its frames a time stamp that is provided, analogously to grid cell codes for space, by timing cells in the entorhinal cortex.

**Fig. 4:**
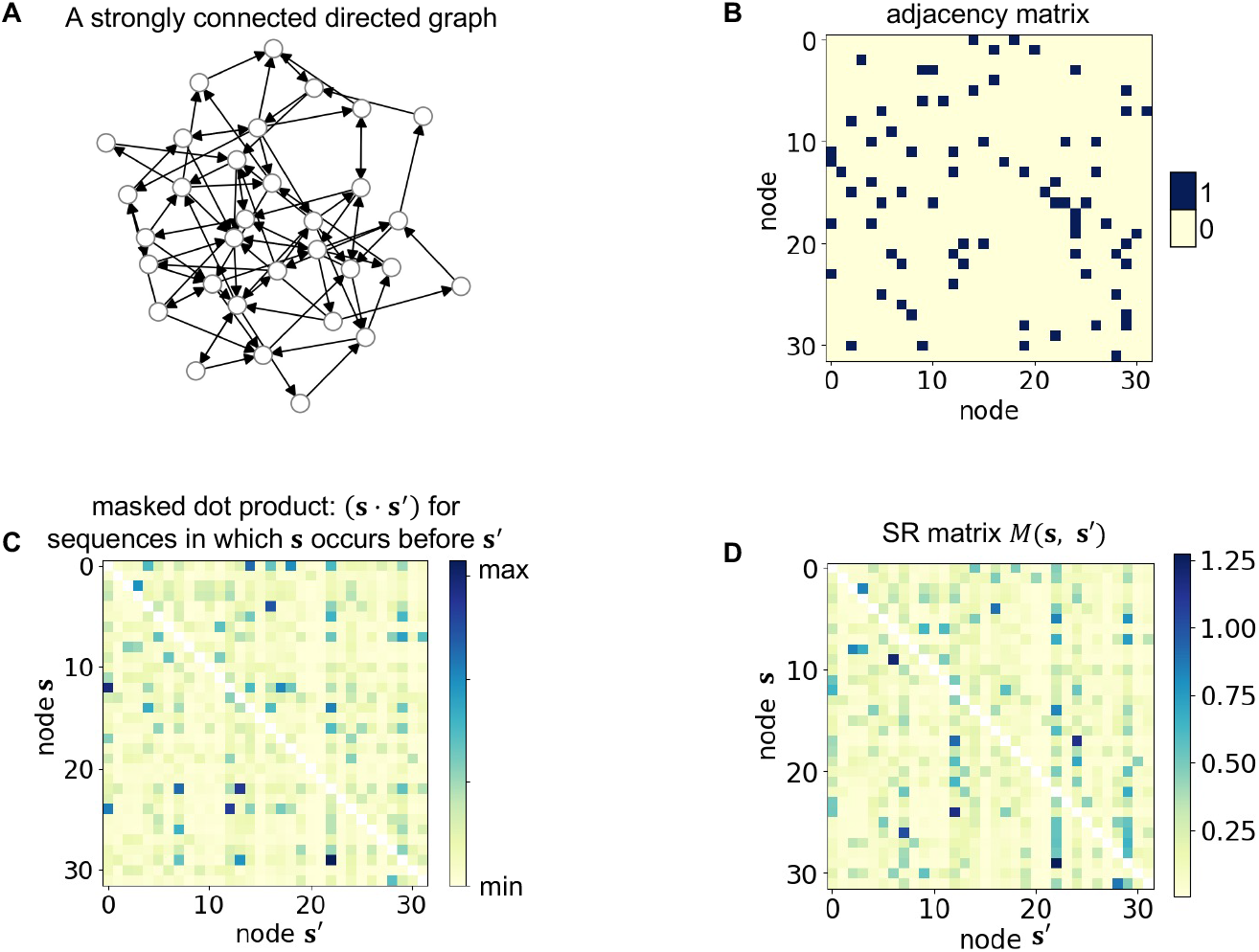
Performance of the CCML for planning in a directed abstract graphs. **(A)** An example strongly connected directed graph with 32 nodes (randomly generated). The proposed method has a near-optimal performance, requiring only 1.92% more steps than Dijkstra, averaged over all start–goal pairs. **(B)** Adjacency matrix of the graph in **A. (C)** Reduced pairwise dot-product heat map of state embeddings. This map accounts for the order of **s** and **s**′ within each trajectory: the reduced embedding of **s**′ retains only those dimensions for which **s** precedes **s**′. **(D)** Successor-representation matrix under a random policy. Note that **C** closely matches **D**.

One should note that the previously described method for planning from the current state **s** = **Q**(**o**) a path to the goal works well only if some path during exploration lead from **s** to the goal. The brain enlarges its repertoire of paths to rewarded locations during replay by joining together several path segments that occurred during exploration (Ou et al., 2025; Pfeiffer and Foster, 2013; Yang et al., 2024). Another method that the brain appears to apply is the creation of episodic memory traces on different temporal and spatial scales. This enables us to recall also episodes where we had visited years ago some far-away location, and steps on the trip to this location on a larger temporal and spatial scale. Hence a multi-scale or hierarchical variant of the CCMl is likely to perform better for problems where also large spatial or temporal distances occur. In any case, a rational default strategy for expanding the CCML if no path is available that leads from the current state to the goal, is to carry out a random walk, or move through some remembered paths until one arrives at states **s**′ for which a path to the goal had already been experienced.

## 3 Discussion

We have shown that the way how episodic memories for sequences of observations are represented in the human hippocampus according to the experimental data of (John et al., 2025; Tacikowski et al., 2024) suggests a simple but clever embedding **Q** of observations into a high-D neural space that can be seen as a cognitive map since it encodes relations between observations in terms of mutual reachability by actions. Therefore this embedding facilitates navigation to any given target observation **o**^∗^. One only needs for that a simple 1-shot learning process for neurons *n*_*a*_ that encode actions **a**. Like for the CML based on prediction learning (Stöckl et al., 2024) this simple architecture and learning process enables online action selection with foresight, expanding the familiar case of navigation in a physical environments where a sense of direction (to the goal) to online navigation in arbitrary concept spaces, or more formally, online generation of shortest paths to given goal nodes in undirected and directed graphs.

The synaptic plasticity mechanism that implements the sequence learning results of (John et al., 2025; Tacikowski et al., 2024) in the human hippocampus are not yet known at present. But the learning window which their data suggests extends over several seconds in both directions from a reference observation, and learning is induced by very few repetitions. These features suggest that the underlying synaptic plasticity mechanism is very similar to the BTSP mechanism that has been identified by (Bittner et al., 2017) in area CA1 of the rodent hippocampus. We have shown that the simple model for BTSP from (Wu and Maass, 2025) suffices for reproducing the emergence of neural codes for sequences of observations in the data from (John et al., 2025; Tacikowski et al., 2024). Experimental data suggest that the LTP window of BTSP is in the range of 6 seconds. This would imply that a neuron can not learn through BTSP to respond to observations in a sequence that stretches over longer time scales. But it well known that different sequences of experiences are integrated into longer sequences during replay, and that the resulting longer sequence is substantially sped up during replay (Ou et al., 2025; Pfeiffer and Foster, 2013). This mechanism could compress also longer sequences into the LTP window of BTSP. Apart from this, the brain apparently also records episodes on a longer time scale, and a corresponding multi-scale extension of the CCML is likely to be able to reproduce that.

Planning and action selection is based in our resulting model on an additional learning mechanism, whereby neurons of that are responsible for initiating specific actions become also responsive to observations of the outcome of this action. Hence these neurons behave after learning like mirror neurons that were discovered over 30 years ago in the primate neocortex (Bonini et al., 2022). Obviously, these mirror neurons play a key role in our CCML model for online planning.

Another interesting feature of the CCML is that it requires a representation of the behavioral goal for online action selection. A number of experimental data have shown that representations of behavioral goals are ubiquitous in the hippocampus, the orbitofrontal cortex, and other brain areas (Basu et al., 2021; Zutshi et al., 2025).

We have shown that the excellent performance of the CCML in online planning of shortest paths in undirected and directed graphs can be analytically understood from the perspective of SR-based RL. SR-based RL provides a powerful method for creating internal data structures that support navigation to any given goal. A number of features of SR-based RL suggest that the human brain employs this method. But a link between this abstract method and the biological implementation level has been missing. In particular, it is not clear how the brain could represent and update a matrix of successor relations, how for a given goal a value function is computed and represented by neurons, and how greedy action selection with regard to this value function can be carried out by neural circuits of the brain. Our CCML model, in conjunction with the recent experimental data on which it is based, provides a simple and biologically plausible neural implementation of SR-based RL.

Apart from this link to SR-based RL the CCML also has a common feature with transformers (Vaswani et al., 2017), since the CCML also learns through self-supervised learning, and takes for action selection the maximum over dot products of target states **s**^∗^ (“queries”) with possible next states that result from currently available actions (“keys”). However, in contrast to transformers, the CCML can already plan well after having experienced just a few trajectories. Curiously, CCML may even rival LLMs on the performance level, since planning capability does not appear to be a strength of current generation LLMs (Momennejad et al., 2023).

Flexible goal-oriented learning capability is also desirable for robots and other edge devices. Because of their energy constraints those methods are most useful in such applications that can be implemented with low energy-consumption in neuromorphic hardware, especially with memristor crossbars for learning and holding the values of synaptic weights. The CCML appears to be highly suitable for such implementations since it only requires simple 1-shot local synaptic plasticity and can be implemented with a small and very simple neural network architecture.

## 4 Methods

### 4.1 Mathematical description

The observation **o**_*i*_ is embedded into the state-space of the CML using the embedding matrix 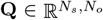:

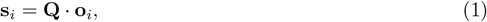

where *N*_*s*_ is the dimensionality of the state space and *N*_*o*_ the dimensionality of the observation **o**_*i*_. This also holds for the target state: **s**^∗^ = **Qo**^∗^.

Structured inputs can be mapped to high-D sparse vectors via BTSP (Wu and Maass, 2025). Here, we use a one-hot vector to represent each resulting observation from an application of the BTSP rule with binary weights: The *j*^th^ node in a graph is represented by **o**_*j*_ = [0, 0, *…*, 1, *…*, 0]^*T*^, which has all zeros except for the *j*^th^ dimension. But one gets similar results if one uses instead biologically realistic sparse and approximately orthogonal input vectors as considered in (Wu and Maass, 2025). During learning, each state neuron becomes selective for all observations within a specific sequence.

This is achieved by setting a state neuron’s incoming weights to 1 for all observation inputs on that sequence:

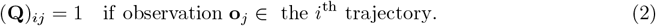

This is equivalent to an application to the simple BTSP rule from (Wu and Maass, 2025) under the assumptions that all these observations arrive within the 6s long LTP window of BTSP and the initial weights have value 0 (Milstein et al., 2021).

Meanwhile, a matrix 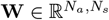 stores incoming weights of action neurons, where *N*_*a*_ is the total number of actions. Each action neuron, corresponding to traversing an edge of the graph in a certain direction, learns to respond to the next state embedding **s**_*i*_ after action **a** is performed. Considering the entire action neuron pool as a long vector, each action is, again, represented by a one-hot code **a**_*j*_ = [0, 0, *…*, 1, *…*, 0]^*T*^, which has all zeros except for its indexed *j*^th^ dimension. The updating rule for **W** follows a Hebbian type:

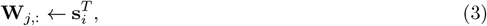

where the post-synaptic **a**_*j*_ activates the action neuron, and selects its incoming weights - the *j*^th^ row of **W**. The rule updates the weights to be the pre-synaptic state **s**_*i*_ resulted from applying action **a**_*j*_. Noting that 1-shot learning with binary synaptic weights, where each action is learned only once, suffices provided one learns **W** after learning **Q**.

During planning, one first computes the current utility values for all actions:

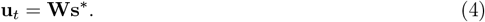

Additional inhibition of actions that cannot be executed in the current state shuts off the corresponding action neurons through inhibition_**s**_(·); a WTA operation then selects the available action with the highest utility value, sets it to 1, and zeros out other action neurons:

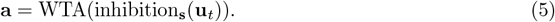

### 4.2 Further details to experiments in Fig. 3

We first focus on the example of a randomly connected graph with 32 nodes, where every node has a random number between two and five edges connecting it to other nodes. The edges of the graph are undirected since they can be traversed in either direction. However, each traversal of an edge in a given direction is viewed as a separate action; thus, there are two actions for every edge.

To explore the graphs, the CCML agents traverse them by taking random walks. The performance depends on two constants: trajectory length *L* and the number of trajectories *N*_*s*_ (which equals the number of state neurons). At each step, the current node along with its previous (*L*− 1) nodes visited during the random walk forms a trajectory. Thus, a new trajectory is generated each time the agent moves. This moving-window approach allows efficient trajectory generation: A training set containing *N*_*s*_ trajectories requires the agent to perform only *N*_*s*_ + *L*− 1 random walk moves.

As a baseline comparison, the Dijkstra graph search algorithm, which computes the shortest path between a pair of nodes, was employed. Our model’s performance is always reported as the percentage increase in the length of each path generated by our model compared to the shortest path length computed by Dijkstra, averaged over all possible (start, goal) nodes.

The initial values for **Q** and **W** were small random values drawn from a Gaussian distribution with (*μ* = 0, *σ* = 10^−4^). Alternatively, these values can be initialized with zeros.

Small world graphs in Fig. 3H can be seen as special challenges, since one needs to identify the precise node that allows escaping from a local cluster. Also, random-walk policy has trouble escaping out of each cluster. Therefore we tested the CML on a graph that had four clusters each consisting of 6 densely interconnected nodes, but only a single connection from one cluster to the next. With the trajectory length *L* = 20 and the number of trajectories *N*_*s*_ = 10^4^, the CCML agent becomes near optimal: it on average takes 4.770 steps as compared to Dijkstra takes 4.714 steps (CCML is 1.19% worse than Dijkstra).

A further concern is that a learning-based online planner may have problems if there are dead ends in a graph: Paths into these dead ends and paths for backing out of them are traversed during learning, and could potentially become engraved into the cognitive map and cause detours when it is applied to planning. To explore how the CCML copes with this difficulty, we designed a graph which contains a large number of dead ends (see Fig. 3I). We found that despite the difficulty of having to deal with dead ends, the CCML managed to achieve the same performance as Dijkstra on this graph with *L* = 10 and *N*_*s*_ = 1000.

The SR matrix in Fig. 3 E captures the structure of the environment based on random walks. For a given adjacency matrix **A** ∈ ℝ^*n*×*n*^ representing the environment’s connectivity (where **A**_*ij*_ = 1 if an edge exists from node *i* to node *j*, otherwise 0), the SR is computed as follows.

We first derive the transition probability matrix **P** ∈ ℝ^*n*×*n*^ of a random walk from the adjacency matrix **A**:

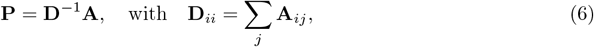

where **D** is the diagonal matrix of node degrees. Specifically, each entry **P**_*ij*_ represents the probability of transitioning from state (node) *i* to state (node) *j* under a uniform random-walk policy.

The SR matrix **M** is then computed analytically as:

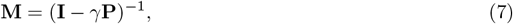

where **I** is the identity matrix, and *γ* ∈ [0, 1) is a discount factor controlling the importance of future states. We set *γ* = 0.8 to obtain the result in Fig. 3E.

### 4.3 Further details to experiments in Fig. 4

We generated strongly connected random directed graphs by first setting a connection probability *p* = 0.15 and then sampling each element in the adjacency matrix according to this probability. We then checked whether the resulting graph was strongly connected; if it was not, we regenerated the graph until a strongly connected graph was obtained. We required strong connectivity here so that a path must exist between any pair of (start, goal) nodes.

The matrix **Q** is initialized as all zeros, and is learned in the same way as in undirected graphs. In addition to recording observations that belong to the same trajectories, we also introduced a “time stamp” for each observation. We used a matrix 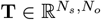 with the same shape as **Q** to record this temporal information:

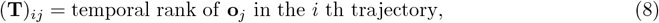

where the temporal rank provide each observation **o**_*j*_ an order of it in the sequence :[1, 2, 3, *…, L*]. If an observation appears more than once in a sequence, we only recorded the temporal rank of its first occurrence.

During planning, the non-zero values in **s**^∗^ are gated depending on the successor state **s**_*i*_ for which we compute utility:

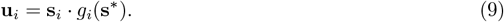

The gating function is computed by comparing two states’ time stamps. Only trajectories where **s**_*i*_occurs before **s**^∗^ are counted:

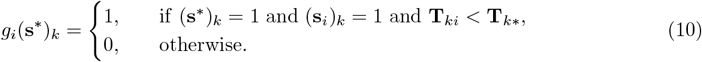

The SR matrix is also computed under random policy as discussed in the previous section. The discount factor *γ* is set to 0.8 to obtain the result in Fig. 4D.

## 5 Acknowledgments

This research was partially supported by the National Science Foundation of the USA (EFRI BRAID project 2318152) and the Austrian Science Fund (FWF) (10.55776/COE12). The authors also gratefully acknowledge the Gauss Centre for Supercomputing e.V. (www.gauss-centre.eu) for funding this project by providing computing time on the GCS Supercomputer JUWELS[1] at Jülich Supercomputing Centre (JSC).

